# Insights from modern & historical genomes of *Neorhizobium vitis*, a new pathogen causing neoplastic growths on grapevine

**DOI:** 10.1101/2025.05.23.655691

**Authors:** Xingyu Wu, Hangwei Xi, Maarten Ryder, Iain R. Searle

## Abstract

Grapevine crown gall is a globally significant disease typically attributed to *Agrobacterium* species carrying tumour-inducing (Ti) plasmids. In this study, we identify *Neorhizobium vitis* as a previously unrecognised crown gall-associated pathogen. Four modern isolates from symptomatic grapevines in South Australia and four historical strains collected in New South Wales in 1989 induced weak but consistent neoplastic growths on sunflower hypocotyls and carrot discs. Long-read genome sequencing revealed that all strains lack canonical Ti plasmids and the oncogenes *iaaM* and *ipt*, which are essential for classical auxin- and cytokinin-mediated tumorigenesis. Phylogenomic analyses reassigned all strains to the *Neorhizobium* clade, correcting the original classification of historical isolates as *Agrobacterium vitis*. The persistence of tumorigenic activity despite the absence of known virulence genes suggests the involvement of alternative, yet uncharacterized pathogenic mechanisms. These findings revise the non-pathogenic status of *Neorhizobium*, establish *N. vitis* as a novel tumorigenic lineage, and underscore the value of re-evaluating legacy isolates with modern genomic tools.

## Introduction

Neoplastic diseases in plants are characterised by abnormal tissue growth, such as tumours or galls, which are often induced by bacterial pathogens. A classical and well-studied example is *Agrobacterium tumefaciens* (Smith and Townsend, 1907), the causal agent of crown gall disease. This pathogen employs a tumour-inducing (Ti) plasmid to transfer oncogenic T-DNA into the plant genome (Larebeke et al., 1974, Chilton et al., 1977, Ooms et al., 1980, Garfinkel and Nester, 1980), reprogramming host cells to overproduce auxin and cytokinin, thereby driving uncontrolled proliferation and gall formation (Klee et al., 1984, Lichtenstein et al., 1984). Related species, such as *Allorhizobium vitis* (formerly *Agrobacterium vitis*), are notorious for grapevine crown gall disease, a globally significant viticultural problem causing vine decline, reduced yields, and premature mortality (Burr et al., 1998).

Beyond Ti plasmid-mediated tumorigenesis, other bacterial mechanisms can induce neoplastic growth in plants. One example is olive knot disease, caused by *Pseudomonas savastanoi* pv. *savastanoi*, which produces auxin and cytokinin through plasmid-encoded biosynthetic genes, triggering hyperplasia without direct DNA integration (Rodr í guez-Palenzuela et al., 2010, Smidt and Kosuge, 1978, Comai and Kosuge, 1980). Another example involves *Pantoea agglomerans*, a typically commensal bacterium that becomes tumorigenic upon acquiring a pathogenicity (pPATH) plasmid carrying a type III secretion system (T3SS) and effector proteins essential for gall formation (Geraffi et al., 2023, Manulis and Barash, 2003). These examples demonstrate that neoplastic plant diseases can result from various bacterial strategies, including plasmid-mediated effector delivery and plant hormone manipulation, underscoring the evolutionary versatility of bacterial pathogenesis.

In contrast, members of the genus *Neorhizobium* separated taxonomically from *Rhizobium* based on multilocus sequence analyses (Mousavi et al., 2014, Sun et al., 2023), are typically non-pathogenic. These Gram-negative soil bacteria form mutualistic symbioses with leguminous plants, establishing nitrogen-fixing root nodules that enhance soil fertility and promote plant growth. Until recently, *Neorhizobium* species were not associated with plant diseases, and their ecological roles were limited to beneficial plant-microbe interactions. However, Haryono et al. (2018) reported a *Neorhizobium* strain NCHU2750 harboring an *Agrobacterium*-type Ti plasmid, demonstrating that even a typically symbiotic bacterium can acquire oncogenic capabilities and induce tumours on plants. This finding suggests that the evolutionary boundary between symbiont and pathogen in the Rhizobiaceae family is more fluid than previously recognised.

Here, we report *Neorhizobium vitis*, a newly identified pathogenic species causing neoplastic growths on grapevines. We identified four *N. vitis* isolates from diseased grapevine plants, whole-genome sequenced the four isolates, performed comparative genomic analyses and tested their pathogenicity on carrot disks and sunflowers. We also sequenced four historical tumorigenic strains isolated by Gillings and Ophel-Keller (1995), corrected their species identification from *A. vitis* to *N. vitis* and showed that these historical isolates are closely related to our recently identified isolates. While grapevine crown gall disease has mainly been attributed to *A. vitis* (Burr et al. 1998), the discovery of *N. vitis* as a causal agent introduces a novel pathogenic lineage within *Neorhizobium*.

## Results & Discussion

To investigate the presence of crown gall-associated bacteria in South Australian vineyards, we collected a symptomatic grapevine exhibiting tumour formation near the graft union and extracted xylem sap using a syringe-based vacuum method (Figure 1, Longchar et al., 2020). The sap was plated on semi-selective Roy and Sasser (RS) medium designed to enrich for *A. vitis* strains (MA, 1983, Burr et al., 1987). We identified fourteen isolates displaying *A. vitis*-like colony morphology and first tested their pathogenicity on sunflower. Of these, four isolates, IRS2293, IRS2294, IRS2295 and IRS2296, induced weak but visible neoplastic growths on sunflower hypocotyls 10–14 days post-inoculation, while the remaining ten isolates produced no neoplastic growth (Figure 2a). Among the IRS isolates, IRS2295 produced the most pronounced tumour phenotype, with visibly larger and more proliferative growths compared to the other strains; however, the tumours remained smaller and less developed than those induced by the positive control strain *A. vitis* K377 (Xi et al., 2021). The weak tumorigenic phenotype suggests these four isolates have low virulence or atypical gall-forming capacity on sunflower.

**Figure 1:**
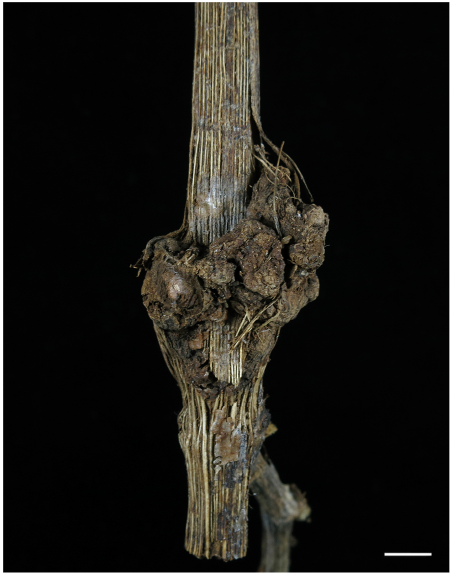
Diseased grapevine plant showing a gall-like growth at the graft union. The plant was collected from the field in 2024. The scale indicates 1.5 cm.

**Figure 2:**
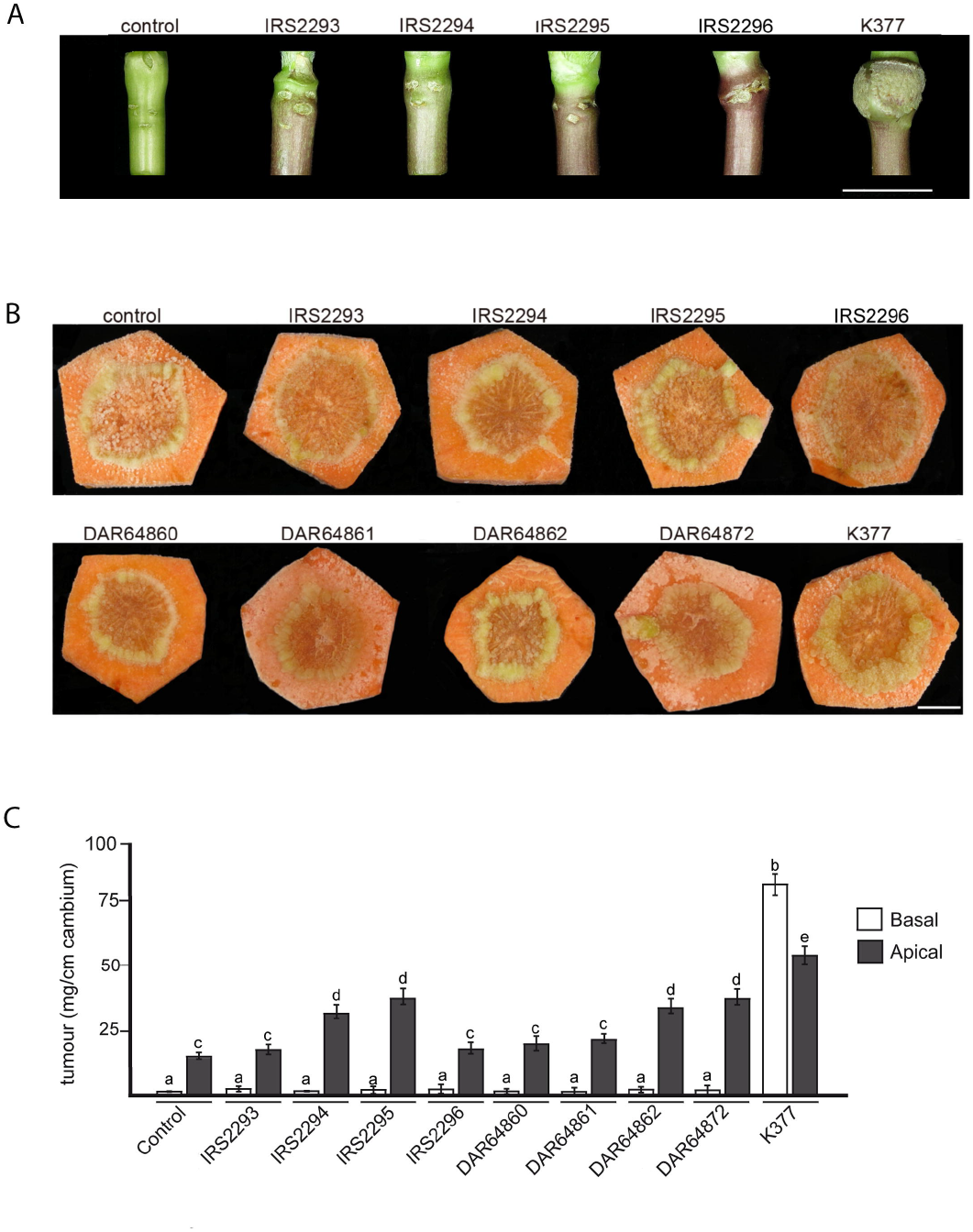
Gall-like growths induced on carrot disks and sunflowers by the strains. **(A)** Sunflower seedling hypocotyls were inoculated with the strains and scored 7 and 14 days later. The image shows sunflowers 14 days after inoculation. The scale indicates 2 cm. **(B)** The apical side of sterile carrot discs were inoculated with candidate bacterial strains to evaluate their gall–inducing potential and scored after four weeks. Tumorigenicity was assessed by the gall weight. Representative images of gall responses are shown. scale indicates 1.5 cm. (**C**) Quantification of the gall mass four weeks after inoculation. 150□µM sodium salicylate was used as a negative control, and *A. vitis* K377 as a positive control. Five replicates were performed of all inoculations. The bar graphs show the mean ± sd with letters showing if there is a significant difference (P<0.05) among the samples as determined by 1-way ANOVA followed by Tukey’s post-hoc test.

We then tested the tumorigenicity of four candidate pathogenic isolates alongside four historically identified tumorigenic strains—K0K1 (DAR64860), K0K2 (DAR64861), K0K3 (DAR64862), and K0K18 (DAR64872)—originally isolated from crown gall-affected grapevines in New South Wales, Australia, in 1989 (Gillings and Ophel-Keller, 1995). The assay was performed using a carrot disc system, a well-established and sensitive method for assessing gall-inducing activity (Wu et al., 2024). Surface-sterilised carrot discs were inoculated with bacterial suspensions and then incubated in the dark under high humidity for four weeks. No tumour formation was observed on the basal surfaces of any carrot discs across all eight tested strains except the well characterized tumourgenic *A. vitis* strain K377 (Figure 2b). However, unexpectedly, weak tumorigenic responses were observed on the apical side of carrot disks based on tumour weight for five strains: IRS2295, IRS2296, K0K2, K0K3, and K0K18 (Figure 2c). Our K377 control produced large apical side tumours. Similar to sunflower, tumour development in these five cases was limited in size, with galls presenting as small, callus-like proliferations in comparison to the virulent strain *A. vitis strain* K377 (Figure 2, Xi et al., 2021). The attenuated gall response contrasts sharply with classical *Agrobacterium fabrum* or *A. vitis* in which robust gall formation typically occurs on both apical and basal surfaces. These findings suggest that the isolates exhibit reduced virulence and potentially rely on different or less efficient mechanisms of host interaction to cause gall formation.

To further investigate the pathogenic mechanisms and phylogenetic relationship to other bacteria of these isolates, we deep-sequenced K0K1 (DAR64860), K0K2 (DAR64861), K0K3 (DAR64862), K0K18 (DAR64872), IRS2293, IRS2294, IRS2295 and IRS2296. Genome assembly was performed using Flye, and gene annotation was conducted with Beav. Our analysis revealed that each isolate possessed two circular chromosomal sequences and between zero and two plasmid sequences (Supplementary Table S1). In the genome assemblies, we searched for tumour induced (Ti) plasmids and unexpectedly did not identify Ti plasmids in any of the eight analysed strains. In stark contrast, we identified the pTi377 plasmid the sequenced, assembly and annotation strain K377.

To determine the phylogenetic placement of these isolates, we constructed a species tree. To do this, we first downloaded protein sequences of two *Rhizobium*, two *Allorhizobium*, two *Agrobacterium* and 41 *Neorhizobium* strains, with one *Pararhizobium* strain as the outgroup (Supplementary Table S2). Protein sequences from the eight isolates, K0K1, K0K2, K0K3, K0K18, IRS2293, IRS2294, IRS2295 and IRS2296, sequenced in this study were added to the dataset. Using OrthoFinder, we identified 1,010 single-copy orthologous genes shared across all samples, performed multiple sequence alignments for each of these single-copy orthologous genes, and gene trees were inferred using IQ-TREE for each gene (Minh et al., 2020). These gene trees were then used to infer a species tree using WASTRAL (Zhang and Mirarab, 2022). Our results showed that the samples from each genus formed distinct monophyletic clades (Figure 3), and the topology of the resulting species tree was consistent with previous studies (Kuzmanović et al., 2022, Wang et al., 2019, Lassalle et al., 2021). The four K0K1, K0K2, K0K3, K0K18 strains that were originally isolated using an *A. vitis* monoclonal antibody were unexpectedly clustered within the *Neorhizobium* clade. This contradicts previous classifications that identified them as *A. vitis* (Gillings and Ophel keller, 1995). Moreover, the four newly isolated strains from this study formed a monophyletic group together with the four K0K1, K0K2, K0K3, K0K18 isolates, indicating a close evolutionary relationship among them.

**Figure 3.**
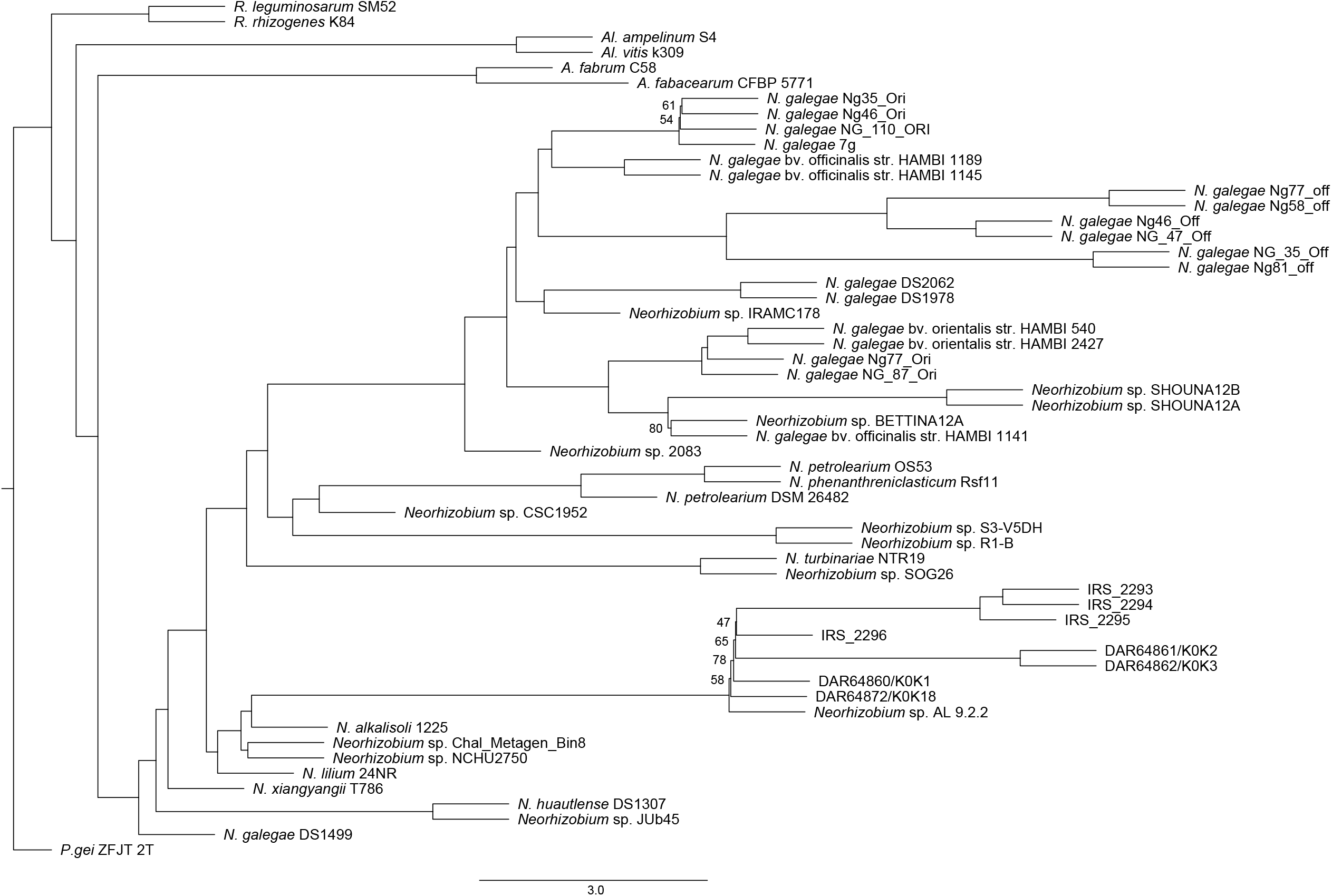
Species tree inferred from 1,010 single-copy orthologous genes. (1000 bootstrap replicates). Branch lengths in coalescent units. Only bootstrap values below 95 are displayed on the tree. The genus names are abbreviated as follows: *A*., *Agrobacterium, Al*., *Allorhizobium, N*., *Neorhizobium, P*., *Pararhizobium, R*., *Rhizobium*.

Virulent tumourgenic *Agrobacterium* and *Allorhizobium* strains are associated with modulating host plant auxin and cytokinin hormones levels by using bacterial encoded, *iaaM* and *ipt* genes, respectively (Spaepen and Vanderleyden, 2011, Kögl et al., 1934, Lichtenstein et al., 1984, Frébort et al., 2011). Unexpectedly, using local blastp searches, we did not identify either *iaaM* or *ipt* genes in the eight K0K1, K0K2, K0K3, K0K18, IRS2293, IRS2294, IRS2295 and IRS 2296 *Neorhizobium* genomes (Supplementary Table S3). Blastp searches for other genes implicated in auxin synthesis, including *iaaH*, and the indole-3-pyruvate pathway genes *iaa1d*, and *ipdC* (Tang et al., 2023), failed to identify any similar genes (Supplementary Table S3). This suggests that these strains increase host plant cell division by another mechanism. Ethylene is known to play tissue and cell-specific roles in plant development, including increased xylem cell division (Abeles et al., 2012, Love et al., 2009), so we then searched the eight genomes for modulators of ethylene levels. However, we did not find any candidate genes, including ACC oxidase in the eight genomes (Supplementary Table S3). In summary, our data support the notation that these strains increase host plant cell division by another unidentified mechanism.

*Neorhizobium* has traditionally been regarded as a non-pathogenic genus within the *Rhizobiaceae*, primarily recognised for forming nitrogen-fixing symbioses with leguminous hosts. However, our findings revise this classification by demonstrating that both newly isolated IRS and historically archived tumorigenic strains are affiliated with *N. vitis*. Specifically, genome sequencing and phylogenomic analysis revealed that four historical grapevine crown gall strains collected in 1989 (K0K1, K0K2, K0K3, and K0K18), previously assumed to belong to the *Agrobacterium* complex, also cluster within the *N. vitis* lineage. Alongside these, newly isolated *Neorhizobium* strains (IRS2293–IRS2296) were found to exhibit weak but consistent neoplastic activity on model hosts, including sunflower hypocotyls and carrot discs.

Despite this tumorigenic phenotype, all *N. vitis* strains examined lacked key molecular hallmarks of classical crown gall pathogens. Whole-genome sequencing confirmed the absence of *Agrobacterium*-type tumour-inducing (Ti) plasmids and the canonical oncogenes *iaaM* and *ipt*, which are central to auxin and cytokinin overproduction and gall development. The lack of these determinants suggests that tumorigenesis in *N. vitis* proceeds through alternative mechanisms. Potential pathways may involve cryptic or novel virulence plasmids, chromosomally encoded genes that influence plant hormone balance, or horizontally acquired effectors that perturb host developmental signalling. The tumorigenic responses observed were generally weak and restricted to specific tissue sites, further suggesting a lower virulence potential or context-dependent pathogenic interaction.

Together, these results redefine *Neorhizobium* as a genus that includes both symbiotic and tumorigenic members and position *N. vitis* as a newly recognized, atypical gall pathogen. The presence of both modern and historical tumorigenic *N. vitis* strains highlights its previously unrecognized contribution to grapevine crown gall disease and underscores the importance of revisiting legacy isolates with modern genomic tools. Further functional and comparative analyses are warranted to uncover the molecular determinants underlying tumorigenesis in this unconventional lineage and to reassess the broader pathogenic potential of other rhizobial genera.

## Materials and Methods

A detailed description of all materials and methods used in this study is available in the Supplementary Methods online. Section 1.1 described the bacteria collection from grapevines; Section 1.2 described the bacteria isolation and culture conditions; Section 1.3 described the pathogenicity tests on the collected bacteria; Section 1.4 described the DNA purification, sequencing, genome assembly and annotation; Section 1.5 described the detail of phylogenetic analysis.

## Data Availability

The authors confirm that all the data used for this study are fully available without restriction. Genome assemblies were deposited in the National Centre for Biotechnology Information (NCBI) under accession numbers listed in the Supplementary Table S2.

## Acknowledgements

This work was supported by a University of Adelaide Waite Research Institute project grant awarded to IRS.

