## Supplemental Data for "Insights from modern & historical genomes of *Neorhizobium vitis*, a new pathogen causing neoplastic growths on grapevine"

**Supplementary data**

### 1 Materials and Methods

#### 1.1 Bacteria collection from grapevines

##### 1.1.1 Source of Vine

Dormant grapevine cuttings (about 40–50 cm in length) exhibiting crown gall symptoms were sourced from vine improvement zones in South Australia in 2024.

##### 1.1.2 Plant growth conditions

The cuttings were surface-sterilised and planted in sterile UC mix (1:1 v/v of sand and peat) under controlled growth chamber conditions at 28 °C with a 16-hour light/8-hour dark photoperiod. The plants were maintained for up to six weeks to promote bud break and shoot development. Successful establishment was defined by the emergence of new stems reaching approximately 10 cm long, which were then used for downstream experimental assays.

##### 1.1.3 Xylem sap collection

Xylem sap was extracted from elongate stems using a syringe-based vacuum method adapted from (Longchar et al., 2020). 4 cm stem segments, 5 cm above the graft union, were collected and surface-sterilised with 70 % ethanol, wiped dry, and trimmed using sterile shears. A P200 pipette tip was fitted onto one end of the stem segment, connected via clear plastic tubing to a 10 ml syringe. The other end of the stem was immersed about 1 cm into 4 ml of sterile phosphate-buffered saline (PBS) in a 5 ml tube. The syringe piston was pulled to create negative pressure and held for 2–4 minutes until liquid uptake ceased. The xylem sap liquid was collected from the syringe for further bacterial isolation.

#### 1.2 Bacterial isolation & culture

##### 1.2.1 Bacteria isolation from grapevine xylem sap

The semi-selective Roy and Sasser-Sodium tartrate (RS-ST) medium was used to isolate Biovar 3 strains. The RS medium is composed of 5.8 g/L Sodium tartrate, 0.14 g/L yeast extract, 0.9 g/L K_2_HPO_4_, 0.7 g/L KH_2_PO_4_, 0.9 g/L MgSO4, 0.2 g/L NaCl, 1.0 g/L boric acid and 15 g/L Agar, pH adjusted to 7.2. After autoclaving and cooling down to 50 ^o^C, 0.08 g/L triphenyl tetrazolium chloride, 0.02 g/L trimethoprim, 0.02 g/L D-cycloserine. 0.02g trimethoprim and 0.25 g cycloheximide were added to the medium before pouring plates. The extracted xylem sap was serially diluted in sterile phosphate-buffered saline (PBS) and plated onto the RS medium. All plates were incubated at 28 °C for 4–7 days until visible colonies developed. Colonies exhibiting morphology consistent with each biovar were sub-cultured for purification and further identification through pathogenicity testing or DNA sequencing.

##### 1.2.2 Bacteria culture conditions

Bacterial isolates recovered from semi-selective media were subsequently cultured on Yeast Extract Mannitol Agar (YMA) or in Yeast Extract Mannitol Broth (YMB), composed of 10 g mannitol, 0.5 g K₂HPO₄, 0.2 g MgSO₄·7H₂O and 1 g yeast extract per litre of distilled water (with 15 g agar added for solid medium), adjusted to pH 6.8–7.0. Agar plates were incubated at 28 °C for 1–3 days until visible colonies developed. Liquid cultures were incubated at 28 °C with shaking on a rotary platform for 1–3 days until reaching the desired optical density (OD).

#### 1.3 Pathogenicity tests

##### 1.3.1 Sunflower stem inoculation assay

The pathogenicity of isolated candidates was first tested using the sunflower stem inoculation assay, adapted from (Stonier, 1969). Sunflower plants (*Helianthus annuus*) were grown under controlled conditions at 25 °C with a 16-hour light/8-hour dark photoperiod until the cotyledon stage (approximately 7–10 days post-sowing). Stems were wounded just below the cotyledon node using three punctures made with a sterile needle. Candidate bacterial strains were cultured overnight and adjusted to ~10⁸ CFU/ml in sterile PBS. A 5 µL aliquot of the bacterial suspension was applied directly to the wound site. Plants were maintained for 2–4 weeks and monitored for gall formation. Tumorigenicity was assessed based on the presence, size, and frequency of gall development compared to mock-inoculated control *A. vitis* K377 strains.

##### 1.3.2  Carrot disc assay

The carrot disc assay was performed following the protocol described by Wu et al. (2024). Candidate bacterial isolates were inoculated onto both the apical and basal surfaces of sterile carrot discs to evaluate their tumorigenic potential. Fresh cultures grown on Yeast Extract Mannitol Agar (YMA) were suspended in 150 µM sodium salicylate to a final concentration of approximately 1 × 10⁷ CFU/ml. A 50 µL aliquot of the bacterial suspension was applied to the cambial ring of each carrot disc. Inoculated discs were incubated in the dark at room temperature for 3–4 weeks. Tumorigenicity was assessed based on the presence, weight of tumour and cambium length at both inoculation sites. Statistical analysis was examined by t-test, 1-way ANOVA with using the GraphPad Prism 9.00 software.

#### 1.4 DNA purification and sequencing

##### 1.4.1 Genomic DNA purification

Bacterial cultures grown in 3 mL of YMB were harvested by centrifugation, and high-quality genomic DNA was extracted using the DNeasy PowerSoil Pro Kit (Qiagen, Germany) according to the manufacturer’s instructions. DNA concentration was measured using a Qubit fluorometer (Thermo Fisher Scientific), and purity was evaluated using a Nanodrop One spectrophotometer (Thermo Fisher Scientific). DNA integrity was confirmed by electrophoresis on a 1% agarose gel.

##### 1.4.2 Nanopore sequencing and genome assembly

400–800 ng of high-quality genomic DNA per sample was used for Nanopore sequencing. Library preparation was performed using the Oxford Nanopore Technologies (ONT) Native Barcode Kit (SQK-NBD.114.24) and required third-party reagents, following the manufacturer’s instructions. Sequencing was carried out on a MinION Mk1B device (ONT, model MIN-114) using an R10.4.1 flow cell (FLO-MIN114). Data acquisition was managed with MinKNOW software (MK1B.23.04.6). Basecalling and demultiplexing were performed using Guppy (v6.5.7) in high-accuracy mode, with reads filtered for a minimum Q-score of 9. Reads were assembled de novo using Flye v2.9.1 with default parameters. The assembled genomes were annotated using Beav v1.4.0 (Jung et al., 2024) in agrobacterium-specific mode (--agrobacterium).

#### 1.5 Phylogenetic tree analysis

We collected protein sequences of two *Allorhizobium*, two *Agrobacterium*, two Rhizobium, 41 *Neorhizobium*, and one *Pararhizobium* strain from the NCBI database, with *Pararhizobium* serving as the outgroup. Protein sequences from eight strains sequenced in this study were also included. Orthogroups were inferred using OrthoFinder v2.5.5 (Emms & Kelly, 2019) in multiple sequence alignment mode (-M msa). Single-copy orthologs shared across all samples were extracted, and their multiple sequence alignments were trimmed using the heuristic method (-automated1) in trimAl v1.5.0 (Capella-Gutierrez et al., 2009). Maximum likelihood (ML) trees were reconstructed for each single-copy ortholog using IQ-TREE v2.4.0 (Minh et al., 2020) with 1000 ultrafast bootstrap replicates (-B 1000). Finally, a species tree was inferred with WASTRAL v1.22.3.7 (Zhang & Mirarab, 2022) based on the ML gene trees, with bootstrap values used to assess branch support (-S).

### 2 Supplementary Data

**Supplementary Table S1**

| **Strain** | **Number, size & conformation of chromosome or megaplasmid** | **Number & size of plasmids** |
| --- | --- | --- |
| K0K1/DAR64860 | 2 chromosomes, 3.51 Mb and 1.04 Mb, both circular | 1 plasmid, 300kb |
| K0K2/DAR64861 | 2 chromosomes, 3.50 Mb and 1.47Mb, all circular | 3 plasmids, 192kb, 91kb and 36kb |
| K0K3/DAR64862 | 2 chromosomes, 3.50 Mb and 1.47Mb, all circular | 2 plasmids, 192kb and 99kb |
| K0K18/DAR64872 | 2 chromosomes, 3.51 Mb and 1.22Mb, all circular | 2 plasmids, 215kb and 95kb |
| IRS2293 | 2 chromosomes, 3.60 Mb and 1.38Mb, all circular | No plasmid |
| IRS2294 | 2 chromosomes, 3.60 Mb and 1.38Mb, all circular | No plasmid |
| IRS2295 | 2 chromosomes, 3.60 Mb and 1.38Mb, all circular | No plasmid |
| *Allorhizobium vitis* K306 | 2 chromosomes, 3.80 Mb and 1.14Mb, all circular | 2 plasmids, 581kb and 263kb |
| *Neorhizobium* sp*.* NCHU2750 | 1 chromosome, 4.32 Mb, circular | 6 plasmids, 765kb, 448kb, 380kb, 222kb, 200kb, 16kb |

**Supplementary Table S2**

| **Strain** | **NCBI accession** |
| --- | --- |
| *Agrobacterium fabacearum* CFBP 5771 | GCA_900039255.1 |
| *Agrobacterium fabrum* str. C58 | GCA_000092025.1 |
| *Allorhizobium ampelinum* S4 | GCA_000016285.1 |
| *Allorhizobium vitis* k309/NCPPB 3554 | GCA_001541345.2 |
| *Neorhizobium alkalisoli* 1225 | GCA_007829835.1 |
| *Neorhizobium galegae* 7g | GCA_021391675.1 |
| *Neorhizobium galegae* bv. officinalis str. HAMBI 1141 | GCA_000731295.1 |
| *Neorhizobium galegae* bv. officinalis str. HAMBI 1145 | GCA_000985915.1 |
| *Neorhizobium galegae* bv. officinalis str. HAMBI 1189 | GCA_000985975.1 |
| *Neorhizobium galegae* bv. orientalis str. HAMBI 2427 | GCA_000986035.1 |
| *Neorhizobium galegae* bv. orientalis str. HAMBI 540 | GCA_000731315.1 |
| *Neorhizobium galegae* DS1499 | GCA_017877135.1 |
| *Neorhizobium galegae* DS1978 | GCA_017877055.1 |
| *Neorhizobium galegae* DS2062 | GCA_030812395.1 |
| *Neorhizobium galegae* NG_110_ORI | GCA_024380075.1 |
| *Neorhizobium galegae* NG_35_Off | GCA_024384505.1 |
| *Neorhizobium galegae* Ng35_Ori | GCA_024384625.1 |
| *Neorhizobium galegae* Ng46_Off | GCA_024384585.1 |
| *Neorhizobium galegae* Ng46_Ori | GCA_024384685.1 |
| *Neorhizobium galegae* NG_47_Off | GCA_023702215.1 |
| *Neorhizobium galegae* Ng58_off | GCA_024384515.1 |
| *Neorhizobium galegae* Ng77_off | GCA_024384545.1 |
| *Neorhizobium galegae* Ng77_Ori | GCA_024384645.1 |
| *Neorhizobium galegae* Ng81_off | GCA_024384605.1 |
| *Neorhizobium galegae* NG_87_Ori | GCA_008806425.1 |
| *Neorhizobium huautlense* DS1307 | GCA_030811685.1 |
| *Neorhizobium lilium* 24NR | GCA_004053875.1 |
| *Neorhizobium petrolearium* DSM 26482 | GCA_017873175.1 |
| *Neorhizobium petrolearium* OS53 | GCA_029854435.1 |
| *Neorhizobium phenanthreniclasticum* Rsf11 | GCA_040066075.1 |
| *Neorhizobium* sp. 2083 | GCA_031455615.1 |
| *Neorhizobium* sp. AL 9.2.2 | GCA_013317225.1 |
| *Neorhizobium* sp. BETTINA12A | GCA_023006485.1 |
| *Neorhizobium* sp. Chal_Metagen_Bin8 | GCA_028283725.1 |
| *Neorhizobium* sp. CSC1952 | GCA_030378245.1 |
| *Neorhizobium* sp. IRAMC:178 | GCA_039631515.1 |
| *Neorhizobium* sp. JUb45 | GCA_004343585.1 |
| *Neorhizobium* sp. NCHU2750 | GCA_003597675.1 |
| *Neorhizobium* sp. R1-B | GCA_004368875.1 |
| *Neorhizobium* sp. S3-V5DH | GCA_004343065.1 |
| *Neorhizobium* sp. SHOUNA12A | GCA_023006505.1 |
| *Neorhizobium* sp. SHOUNA12B | GCA_023006485.1 |
| *Neorhizobium* sp. SOG26 | GCA_003491345.1 |
| *Neorhizobium turbinariae* NTR19 | GCA_023223505.1 |
| *Neorhizobium xiangyangii* T786 | GCA_020531965.1 |
| *Pararhizobium gei* ZFJT.2T | GCA_029223885.1 |
| *Rhizobium leguminosarum* SM52 | GCA_004306555.1 |
| *Rhizobium rhizogenes* K84 | GCA_000016265.1 |
| DAR64860/K0k1 | CP191458-CP191460 |
| DAR64861/K0K2 | CP191453-CP191457 |
| DAR64862/K0K3 | CP191449-CP191452 |
| DAR64872/K0K18 | CP191445-CP191448 |
| IRS_2293 | CP191441-CP191442 |
| IRS_2294 | CP191439-CP191440 |
| IRS_2295 | CP191443-CP191444 |

**Supplementary Table S3**

| **Samples** | ***ACO1^a^*** | ***ACO2^b^*** | ***IaaH^c^*** | ***IaaM^d^*** | ***IaaL^e^*** | ***ipt^f^*** | ***ipdC^g^*** | ***pehA^h^*** |
| --- | --- | --- | --- | --- | --- | --- | --- | --- |
| IRS_2293 | N | N | N | N | N | N | N | N |
| IRS_2294 | N | N | N | N | N | N | N | N |
| IRS_2295 | N | N | N | N | N | N | N | N |
| IRS_2296 | N | N | N | N | N | N | N | N |
| K0K1/DAR64860 | N | N | N | N | N | N | N | N |
| K0K2/DAR64861 | N | N | N | N | N | N | N | N |
| K0K3/DAR64862 | N | N | N | N | N | N | N | N |
| K0K18/DAR64872 | N | N | N | N | N | N | N | N |
| *A. fabrum* C58 | N | N | P | P | N | P | N | N |
| *Al. vitis* K306 | N | N | P | P | N | P | N | P |
| *Ps. savastanoi* pv. savastanoi NCPPB 3335 | N | N | P | P | P | N | N | P |

N not detected, P present. The threshold of positive is amino acid identity > 30% and query coverage > 70%.

Details of query sequences for blast search:

***^a^*** *ACO1*: *ACO1* from *Ar. thaliana* (NCBI Reference Sequence: NP_179549.1).

***^b^*** *ACO2*: *ACO2* from *Ar. thaliana* (NCBI Reference Sequence: NP_176428.1).

***^c^*** *iaaH*: *iaaH* from *Al. vitis* (NCBI Reference Sequence: WP_070167543.1), *A. fabrum* C58 (WP_010974823.1), *A. tumefaciens* Ach5 (WP_010892363.1), *Al. ampelinum* S4 (WP_012649066.1), *Ps. savastanoi* pv. savastanoi NCPPB 3335 (WP_002556047.1).

***^d^*** *iaaM*: *iaaM* from *Al. vitis* k377 (NCBI Reference Sequence: WP_032494182.1), *A. fabrum* C58 (WP_010974824.1), *A. larrymoorei* FPH_AR2 (WP_081307295.1), *A. rubi* A19/93 (WP_010891461.1), *A. tumefaciens* Ach5 (WP_162163087.1), *A. tumefaciens* Q15/94 (GenBank: QTG17272.1), *Al. vitis* CG142 (WP_081089077.1), *Al. ampelinum* S4 (WP_324611384.1), *R. rhizogenes* CA84 (WP_174045210.1).

***^e^*** *iaaL*: *iaaL* from *Ps. savastanoi* pv. savastanoi NCPPB 3335 (UniProtKB/Swiss-Prot: P18204.1).

***^f^*** *ipt*: *ipt* from *A. larrymoorei* FPH-AR2 (NCBI Reference Sequence: WP_065657522.1), *A. tumefaciens* Q15//94 (WP_333722764.1), *A. vitis* CG142 (WP_032488312.1), *A. fabrum* C58 (WP_010891460.1), *A. rubi* A19/93 (WP_065698664.1), *A. tumefaciens* Ach5 (WP_010892365.1), *Al. vitis* K377 (WP_070167542.1), *R. rhizogenes* CA84/95 (WP_174045225.1), *Al. ampelinum* S4 (WP_012649069.1).

***^g^*** *ipdC*: *ipdC* from *Pa. agglomerans* FDAARGOS 1447 (NCBI Reference Sequence: NZ_CP077366.1), *Ps. putida* (GenBank: OLS60705.1)

***^h^*** *PehA*: PehA from *R. laguerreae* TLR6 (NCBI gene ID 67488026) and *Al. vitis* (Uniprot accession: P77818)

*A*., *Agrobacterium*, *Al*., *Allorhizobium*, *Ar*., *Arabidopsis*, *Pa*., *Pantoea*, *Ps*. *Pseudomonas, R*., *Rhizobium*.
